# Detection of hematopoietic stem cell transcriptome in human fetal kidneys and kidney organoids derived from human induced pluripotent stem cells (iPSC)

**DOI:** 10.1101/2021.01.22.427745

**Authors:** Jin Wook Hwang, Christophe Desterke, Julien Loisel-Duwattez, Frank Griscelli, Annelise Bennaceur-Griscelli, Ali G Turhan

## Abstract

**Background:** In mammalians, hematopoietic stem cells (HSC) arise in the dorsal aorta from the hemogenic endothelium, followed by their migration to fetal liver and to bone marrow. In zebrafish, kidney is the site of primary hematopoiesis. In humans, the presence of HSC in the fetal or adult kidney has not been established.

**Methods:** We analyzed the presence of HSC markers in human fetal kidneys by analysis of single-cell datasets. We then analyzed in kidney organoids derived from iPSC, the presence of hematopoietic markers using transcriptome analyses.

**Results:** 12 clusters were identified of stromal, endothelial, and nephron cell type-specific markers in the two fetal stage (17 weeks) kidney datasets. Among these, expression of hematopoietic cells in Cluster 9 showed expression of primitive markers. Moreover, whole transcriptome analysis of our iPSC-derived kidney organoids revealed induction of the primitive hematopoietic transcription factor RUNX1 as found in the human fetal kidney cortex.

**Conclusions:** These finding support the presence of cells expressing HSC transcriptome in human kidney. The mechanisms of the appearance of the cells with the same transcriptional features during iPSC-derived kidney organoid generation requires further investigation.

## Introduction

Hematopoietic stem cells (HSC) are characterized by their capacity of both self-renewal and differentiation into blood and immune cell lineages throughout the life of the individual in a stem cell-regulating microenvironment, or HSC niche. HSC-niche interactions in bone marrow, liver and kidney have been extensively studied using vertebrate animal models, including mice, frogs, zebrafish, and chickens [1,2]. During mammalian hematopoiesis, the most primitive hematopoietic cells migrate from the aorta–gonad–mesonephros (AGM) region to the fetal liver and to the bone marrow which is the site of adult hematopoiesis [1]. However, the persistence of some degree of hematopoietic activity in adult tissues is possible as this has been suggested by the discovery of donor-derived chimeric hematopoiesis after liver transplantation showing the contribution of donor-derived cells to hematopoiesis [3]. To our knowledge, there has been no study analyzing the possibility of donor-derived hematopoiesis after kidney transplantation. It should be reminded that in the majority of cases, kidney transplants are performed using kidneys from deceased donors [4,5]. HSC-kidney niche interactions have been studied in many reports. For example, zebrafish kidney stromal cell lines can support and maintain early hematopoietic precursors and differentiation of lymphoid, myeloid, and erythroid precursors [4].

Recent advances in single-cell RNA sequencing technology are leading to new discoveries and validation in fetal organs and organoids [5–7]. Here, we first analyzed the presence of transcriptional markers of HSC in fetal kidneys through analysis of a single-cell dataset described by Lindström NO et al [5–7]. We then performed a transcriptome analysis of iPSC-derived kidney organoids. We show that HSC-related markers can be detected in both in fetal kidneys and the human iPSC-derived kidney organoids.

## Results

### Human fetal kidney cortex harbors cells expressing hematopoietic transcripts

Single cell transcriptome is a powerful technology to investigate cell heterogeneity in a tissue. Lindström NO et al [5–7] performed these experiments in cortexes tissues isolated from two human fetal kidneys (17 weeks) by 10X genomics technology. This work allowed us to perform in silico analyses. To this end, we merged and analyzed with Seurat package the 2 respective MTX files generated by cell ranger in order to suppress batch error and performed downstream unsupervised analysis. To build the common matrix of the two samples, genes which were found expressed in minimum 5 cells by sample were conserved. After merging of the data from the two kidney samples, the Seurat digital matrix comprised 7860 cells for 18119 transcripts. During batch correction with canonical correlation, we observed that the 2 kidney samples were found well superposed in first factorial map of canonical correlation (Fig. S1a) and the shared correlation strength decrease on the thirty components of canonical correlation (Fig. S1b). tSNE analysis on the common variable genes on the 40 principal components of the principal component analysis allowed to identify 12 clusters (Fig. 1a) reproducible in both kidneys (Fig. S1c). Major of the tSNE central cells comprising clusters 3,2,0,1 expressed mesoderm transcription factor TCF21 (Fig. 1a and Fig. S2), also expression of TCF21 is positive in cells from cluster 6 which highly expressed matrix molecules such as Lumican (LUM) (Fig. 1a and Fig. S2) Decorin (DCN) and Collagens (COL3A1, COL1A1, COL1A2) (Fig. 1b). In cluster 4, cells were found to be positive for KDR (Fig. 1a) and CD34 (data not shown), suggesting an expression profile corresponding to endothelial cells. Cells identified in cluster 7 have a high expression of downstream NOTCH pathway transcription factor HEY1 such as cells from cluster 0 which are central proximal from this position during tSNE analysis (Fig. 1a). Some cluster of cells which are left eccentric (clusters 8 and 11) expressed tubular markers such as FXYD2 (Fig. 1a and Fig. S2) encoding the sodium/potassium-transporting ATPase subunit gamma. Cluster of cells number 10, also left eccentric during tSNE analysis expressed some podocyte markers such as PTPRO: Protein Tyrosine Phosphatase, Receptor Type O (Fig. 1a and Fig. S2), but also SOST: Sclerostin and PODXL: podocalyxin like (Fig. 1b). Cluster of cells number 5 expressed specifically the renin molecule a well-known renal molecule (Fig. 1a and Fig. S2). These results suggest that tSNE analysis performed post canonical analysis in these 2 merged samples reflect the cell diversity compatible with kidney organ at this stage of development described in the original paper [8]. Surprisingly, in unsupervised tSNE analysis of human kidney cortex, we found the left-top eccentric cluster number 9 (Fig. 1a) which is principally defined by the specific expression of SRGN (Fig. 1a, Fig. 2a and Table S1). Serglycin (SRGN) is known to be a hematopoietic cell granule proteoglycan. In this cluster of cells, there is also a specific expression of hematopoietic cluster of differentiation such as PTPRC alias CD45 and CD44 (Fig. 2b). Some molecules such as CD74 and HLA-DRA implicated in antigen presenting cells functionalities are also expressed in this cluster of cells (Fig. 2c). Interestingly, a fraction of cells from cluster 9 also expressed primitive hematopoietic transcription factors such as SPI1 (alias PU.1) and RUNX1 (Fig. 2d). Some of the cells from the same cluster also expressed CXCR4 receptor which is well known to be expressed on primitive human hematopoietic cells for their homing function. This original results on cluster 9 suggest the presence of cells with hematopoietic transcriptome with some of them expressing primitive markers in the human kidney cortex at fetal stage (17 weeks).

**Fig. 1.**
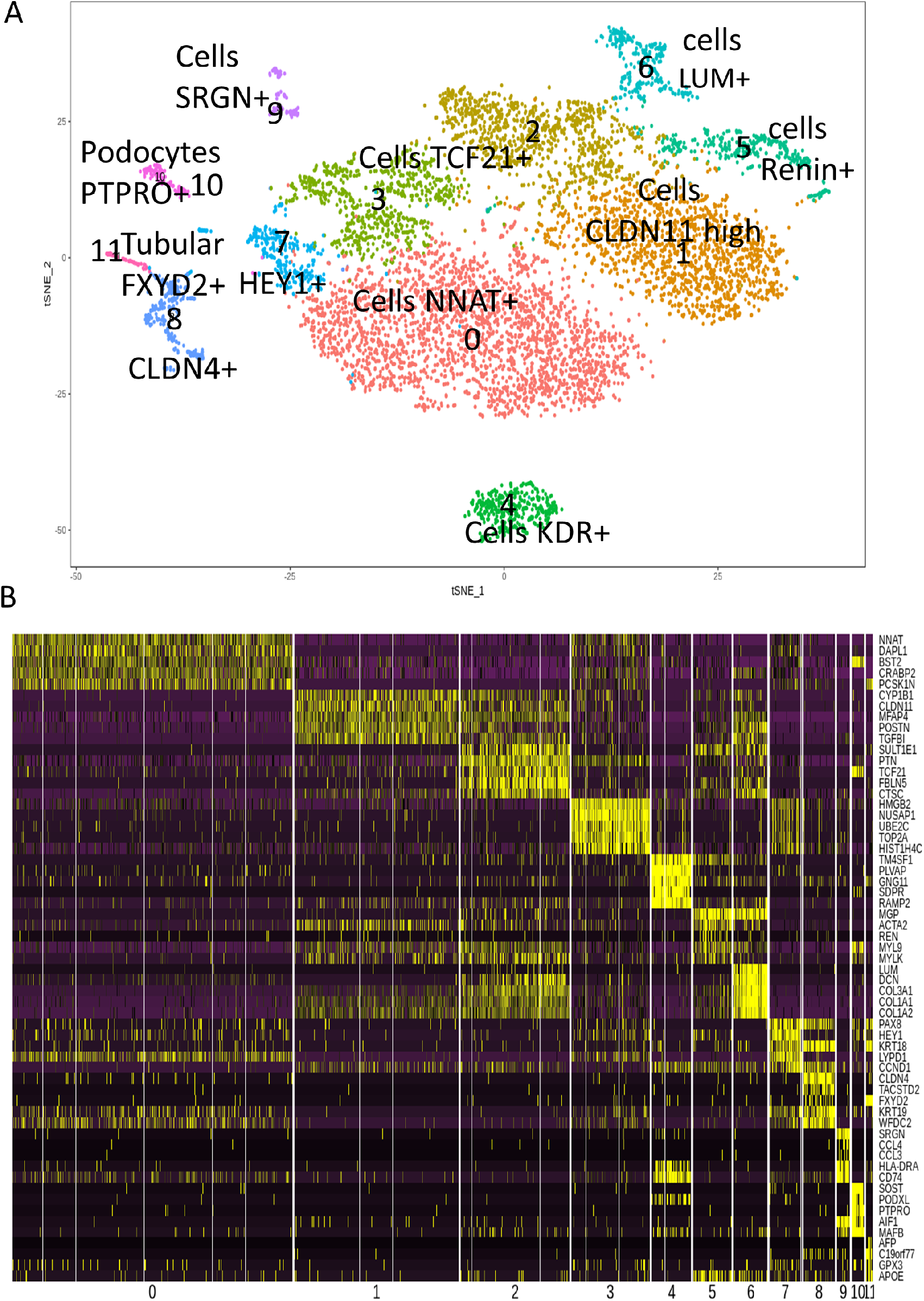
Cell heterogeneity in human fetal kidney cortex by single cell transcriptome. **a** tSNE plot with eleven cell clusters from the combined analysis of the merged fetal kidney cortex (2 human kidneys 17 weeks; 7860 cells) after canonical correlation. **b** Heatmap with the expression pattern of the top five cluster-specific genes in 11 clusters identified in human fetal kidney cortex.

**Fig. 2.**
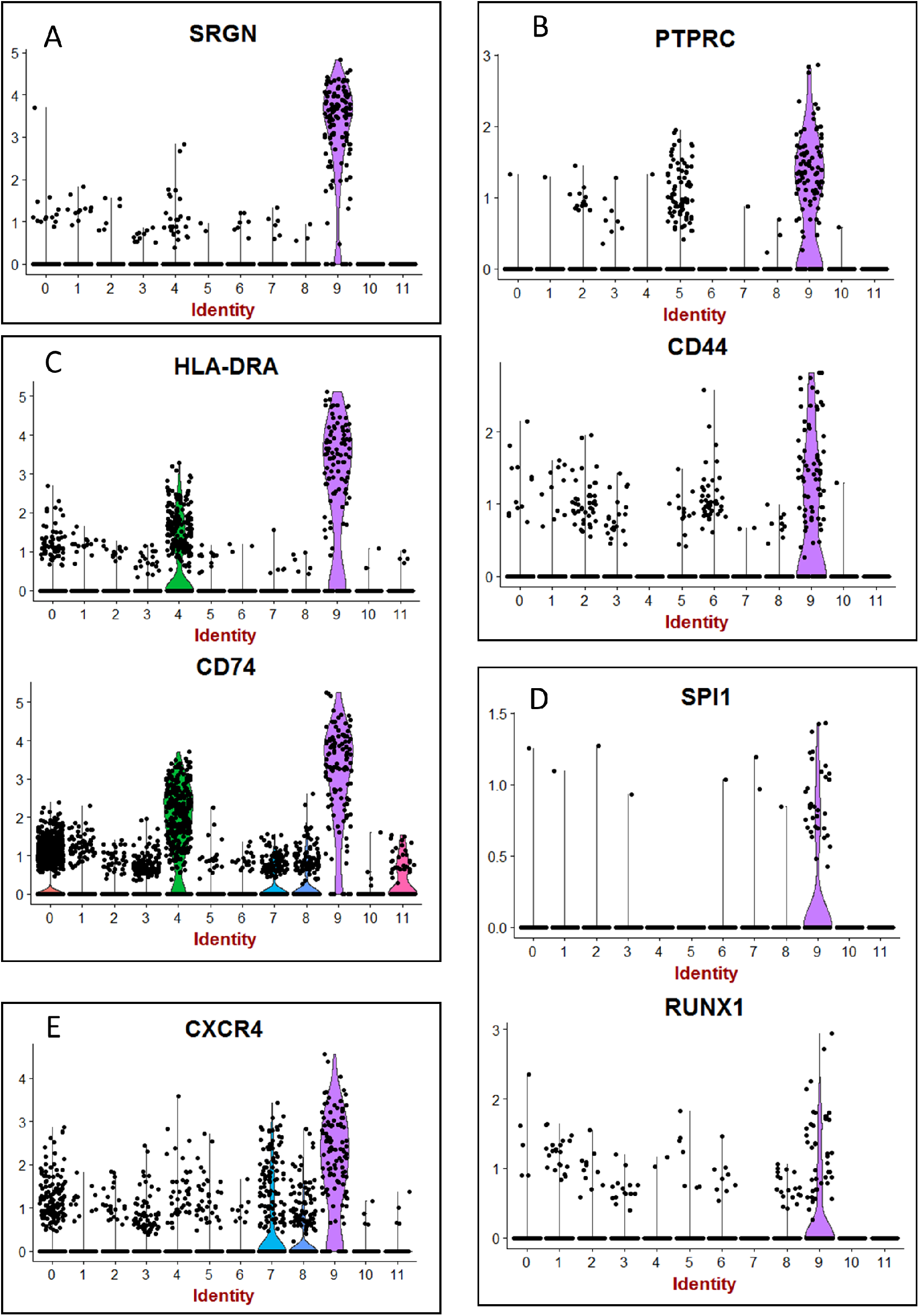
Hematopoietic transcripts detected in human fetal kidney cortex by single cell RNA sequencing. **a** Violinplot of SRGN expression. **b** Violinplot of hematopoietic clusters of differentiation (PTPRC Alias CD45). **c** Violinplot of expression of transcripts of differentiated hematopoietic cells (HLA-DRA: HLA-DR Alpha). **d** Violinplot of expression of SPI1 (PU. 1) and RUNX1. **e** Violinplot of expression for CXCR4 receptor.

### Generation and characterization of iPSC-derived kidney organoids

Human iPSC-derived kidney organoids have been generated as previously described [9]. Briefly, iPSC aggregates were generated in E8 media and Geltrex matrix leading to spontaneous formation of complex kidney organoids at days 12-14 of the culture. (Fig. 3a). We characterized iPSC-derived kidney organoids using whole-mounting staining with confocal imaging. As can be seen in Fig. 3b, glomeruli-like structures, which contained cells that stained for the nephron marker Nephrin were easily identified [10]. Moreover, ultrastructure analyses revealed cell-cell junctions and the podocyte foot process formation in kidney organoids (Fig. 3c).

**Fig. 3.**
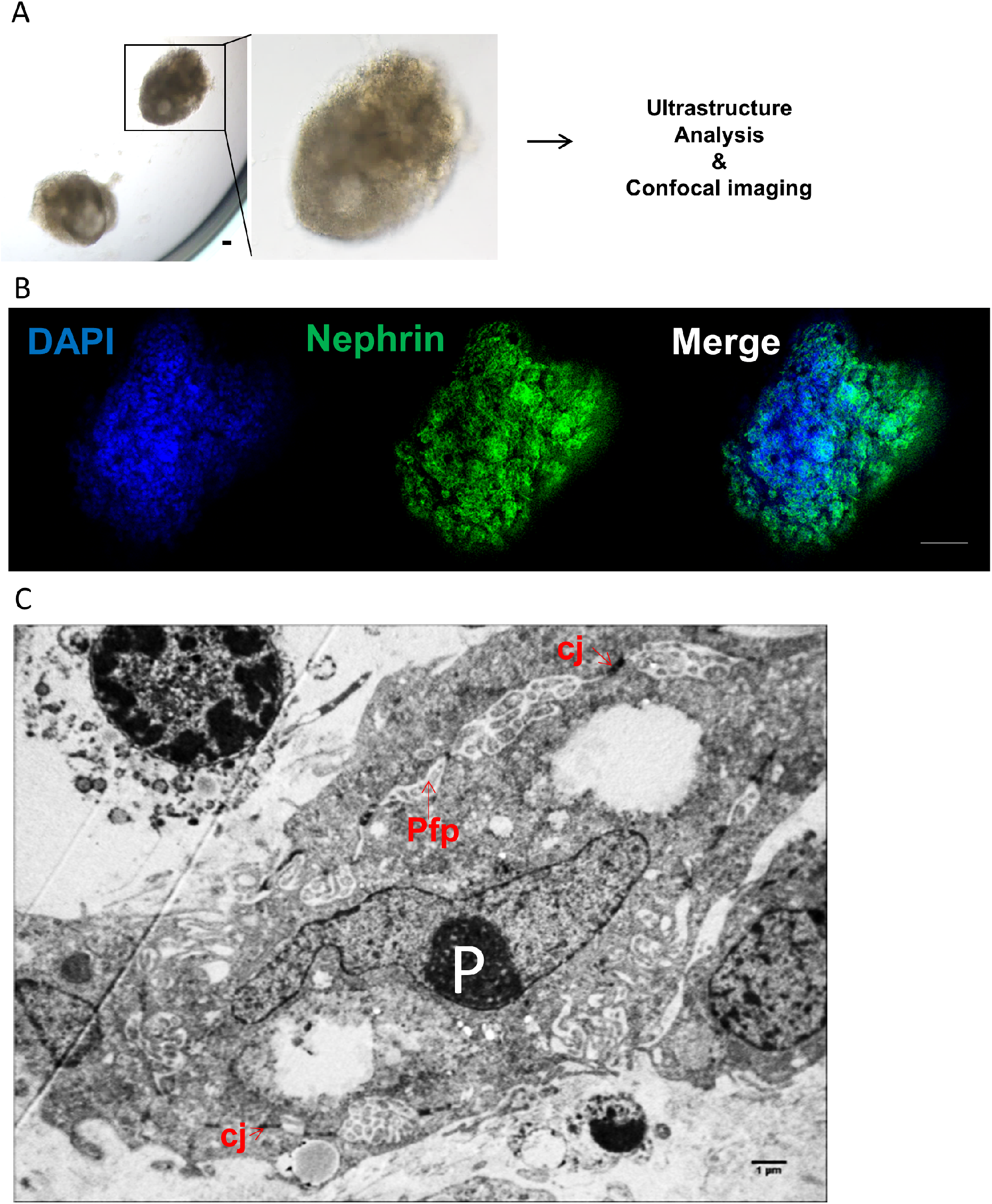
Characterization of human iPSC-derived kidney organoids. **a** Optical image of iPSC-derived kidney organoids at day+14. Scale bar: 100 μm. **b** Confocal analysis and whole-mount staining for Nephrin in iPSC-derived kidney organoids showing nephron vesicles. Scale bar: 50 μm. **c** Representative electron microscopy image of podocytes in iPSC-derived kidney organoid at day+ 14 showing podocytes (P), podocyte foot process (Pfp) and cell-cell junctions (cj). Scale bar: 1 μm.

### Detection of a hematopoietic transcriptome program iPSC-derived kidney organoids

We performed, in duplicate, whole transcriptome analysis of iPSC derived kidney organoids as compared to native iPSC with Clariom S human technology. After RMA normalization, we identified 3546 differentially expressed genes (DEG) with LIMMA algorithm (Fig. 4a) comprising 1432 up-regulated genes. This DEG profile allowed to discriminate experimental sample groups by unsupervised classification (Fig. 4b). After functional enrichment on WikiPathway database, we identified a hematopoietic program in iPSC-derived kidney organoids. Especially, we uncovered an up regulation of RUNX1 and CD34 corresonding to genes expressed in hematopoietic stem cells and that of FLI1, CXCR4, MXI1 downstream at erythrocyte and megakaryocyte progenitor levels. There was also a repression of MYB megakaryocytic repressor (Fig. 4c). These results suggest the implication of a hematopoietic transcriptional program in our iPSC-derived kidney organoids and especially induction of the primitive hematopoietic transcription factor RUNX1 which was also detectable at the single cell level (Fig. 2d) in human ex vivo fetal kidney cortex.

**Fig. 4.**
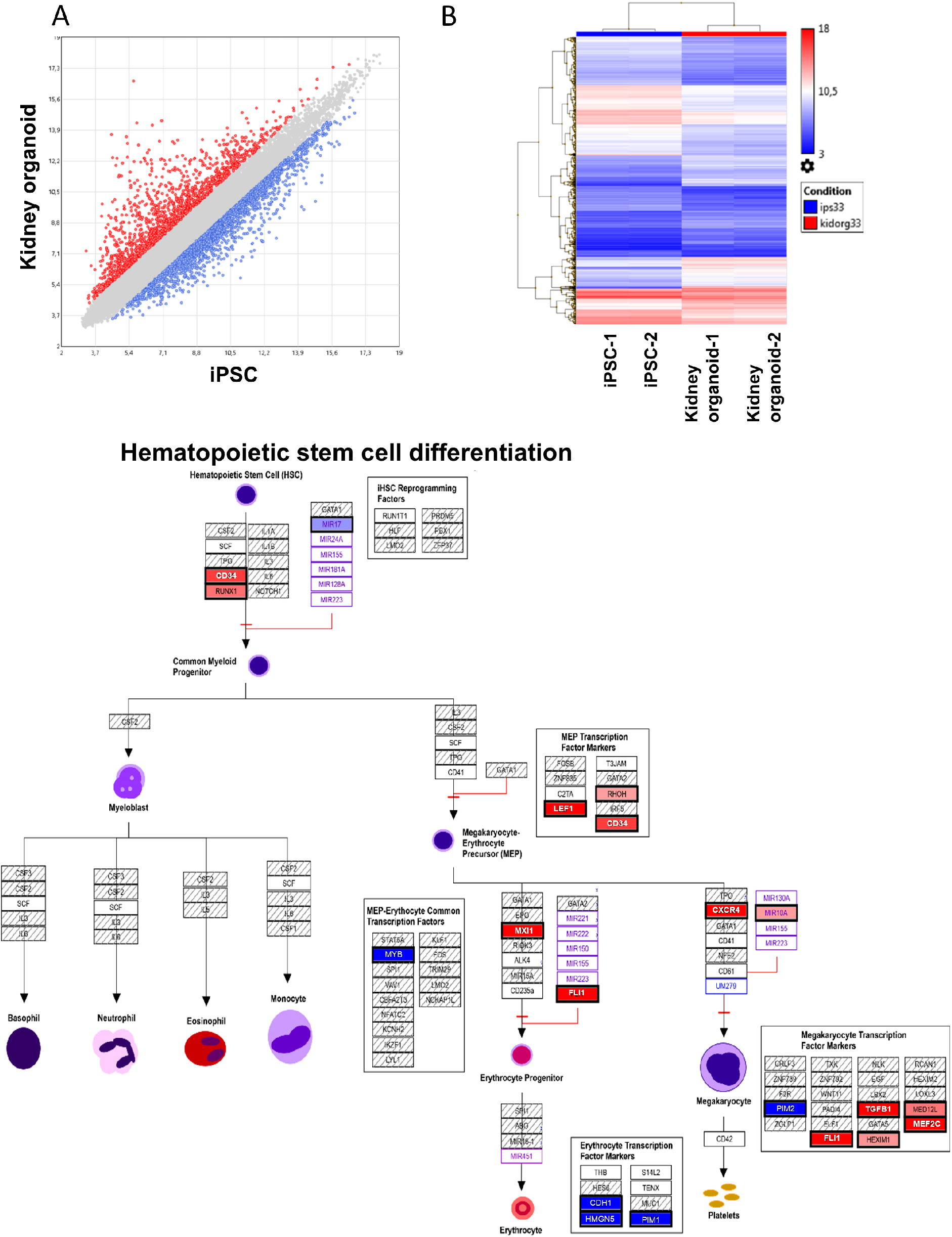
Transcriptional program induced in kidney organoid derived from human iPSC. **a** Scatterplot of differential expressed genes found in transcriptome of kidney organoid versus iPSC. **b** Expression heatmap with unsupervised classification performed on differentially expressed genes induced during kidney organoid differentiation from iPSC. **c** Functional enrichment performed on WikiPathway database showing potential implication hematopoietic stem cell function during differentiation of kidney organoid derived from human iPSC.

## Discussion

The involvement of kidney in hematopoiesis has been clearly demonstrated in zebrafish [11]. In humans, the most primitive hematopoietic cells arise from mesodermal lineage in AGM through hemogenic endothelium [12]. Kidney is also a tissue developed from mesoderm but the presence of cells with HSC transcriptome has not been studied. Here, we first analyzed the HSC markers in fetal kidney through analysis of fetal kidney single cell dataset analyses. In human fetal kidney cortex, we found some cells expressing RUNX1 in a cluster of cells which harbored expression hematopoietic markers (Cluster 9 on Fig. 1a and 1b). In cluster 9 of human fetal kidney sc-RNAseq, high expression of hematopoietic markers was confirmed by the presence of Serglycin (SRGN)-positive cells (Fig. 2a) as well as cells expressing PTPRC alias CD45 Leukocyte Common Antigen and CD44 (receptor of hyaluronic acid) (Fig. 2b). Serglycin (alias hematopoietic proteoglycan core protein) is a protein found in secretory granules of myeloid cells as well as in platelets. Our analysis showed also the presence of cells positive for MHC class II molecule HLA-DRA and CD74 (Fig. 2c) and most interestingly, cells expressing of hematopoietic transcription factors SPI1 and RUNX1 (Fig. 2d). Finally, in this cluster 9 of fetal kidney we found a higher expression of CXCR4 receptor of CXCL12 implicated in migration properties of HSCs (Fig. 2e). All these results allowed to suggest the presence of cells harboring hematopoietic transcriptome in human fetal kidney.

We then analyzed the transcriptome of iPSC-derived kidney organoids and performed differential expression analysis of the kidney organoid versus parental iPSCs. Microarray analysis revealed important regulation of transcriptional program of these cells during their differentiation (Fig. 4a). This differentially expressed program allowed to discriminate group samples by unsupervised classification (Fig. 4b). After functional enrichment performed on up-regulated genes during the differentiation process of the iPSC-derived kidney organoids, we observed the induction of HSC markers such as RUNX1 and CD34 (Fig. 4c). It is well established that RUNX1 along with a cis-regulatory elements integrating the GATA, ETS, and SCL transcriptional networks, plays a major role in HSC generation [13]. We also found induction of FLI1 during the differentiation of iPSC-derived kidney organoid. SPI1 (alias PU.1), the main target downstream RUNX1 [14] is also a master regulator of hematopoiesis as it prevents excessive HSC division and exhaustion by controlling the transcription of multiple cell-cycle regulators [15]. Association of SPI1 and RUNX1 are comprised in a combination of 7 transcription factors which are sufficient to convert hemogenic endothelium into hematopoietic stem and progenitor cells that engraft myeloid, B and T cells in primary and secondary mouse recipients [16]. In transcriptome analyses of iPSC-derived kidney organoids, we found an up-regulation of MYB which is known to participate to cell fate decisions between erythropoiesis and megakaryopoiesis in human hematopoiesis [17]. Amongst the hematopoietic transcripts identified in human fetal kidney cortex, we have also detected the expression of which, CXCR4 in relation with its ligand CXCL12, is involved in homing of hematopoietic cells to the bone marrow [18].

Our data has some limitations including the fact we can not exclude the presence of mesodermal cells undergoing the fate of hematopoietic differentiation during our kidney organoid differentiation. Secondly, we could not identify the presence of cells with HSC functionality (self-renewal; differentiation) in the current experiments. However these data suggest that at some point during embryonic development, a special “kidney niche” could appear transiently in humans. The identification of such a niche or its molecular counterparts could be of major interest to amplify human HSC for transplantation purposes, such has been described in zebrafish [4,19]. It is known that zebrafish embryonic stromal trunk (ZEST) cells derived from the HSC emergence site are functionally similar to the mammalian AGM niche cells. Moreover, ZEST cells and kidney cell lines have similar signaling properties. [19,20]. Our results suggest that a “kidney microenvironmental niche” niche could be of interest to generate conditions for HSC culture and expansion.

## Materials and Methods

- KEY RESOURCES TABLE

| REAGENT or RESOURCE | SOURCE | IDENTIFIER |
| --- | --- | --- |
| Antibodies |  |  |
| Nephrin | Abcam | ab85379 |
| DAPI | Sigma-aldrich | D9542 |
| **Chemicals, Peptides, and Recombinant Proteins** |  |  |
| Essential 8 basal medium | Thermo Fisher Scientific | A1516901 |
| Essential 8 supplement | Thermo Fisher Scientific | A1517101 |
| Geltrex® LDEV-Free Reduced Growth Factor Basement Membrane Matrix | Thermo Fisher Scientific | A1413202 |
| ROCK inhibitor | Global stem | GSR-6102 |
| **REAGENT or RESOURCE** | **SOURCE** | **IDENTIFIER** |
| Experimental Models : Cell Lines |  |  |
| Human iPSC: PB33 | Human | - |
| Osmium tetroxide solution | Sigma-aldrich | 75632 |
| Glutaraldehyde grade I | Sigma-aldrich | G5882 |
| **Software and Algorithms** |  |  |
| ImageJ |  |  |

### Generation of iPSC

The iPSC line used is this study was generated using Sendaï virus-mediated gene transfer of the four “Yamanaka” factors as previously described (9).

### Generation of Kidney organoids

iPSCs were maintained on Geltrex (Stem Cell Technologies, Inc) coated flat culture dish in E8 media (Stem Cell Technologies, Inc) according to manufacturer’s guidelines. Colonies were manually harvested at 60-80% confluence. Cells were then collected and dissociated into single cells using EDTA. Cells (1×10^6 or 1×10^5/well) were put onto ultra low attachment 24 well or 96 well plate (Corning, Inc) to allow them to form aggregated in suspension with ROCK inhibitor (2-5 μmol). Cell aggregates were cultured in E8 medium (Stem Cell Technologies) with daily medium change for 6-7 days. Control iPSC-A (iPSC-aggregates) were plated on a Geltrex (Stem Cell Technologies) in 96 well plate or 8 well culture chamber. And then aggregates were treated E8 medium (Stem Cell Technologies) with daily medium change for 12-14 days. Images were taken using a NIKON microscope.

### Whole-mount immunostaining of 3D kidney organoids

Kidney organoids cultured on 96-well culture dishes were washed with phosphate-buffered saline (PBS), fixed with 4°C paraformaldehyde in PBS for 120 min, permeabilized with 0.2% Triton X-100 (Sigma) in PBS and blocked in 10% serum. For nephrin staining, the antibody (Nephrin (Cat#ab85379; Abcam) was diluted in PBS containing 10% serum and washed in PBS. Samples were incubated with secondary antibodies in antibody dilution buffer, then washed in PBS. Nuclei were labeled with DAPI mounting medium. Visualization and capture were realized with a Zeiss confocal microscope.

### Transmission Electron Microscopy (TEM)

Kidney organoids were gently centrifuged, and pelleted before the TEM process as follows. The cells were fixed in 2.5% glutaraldehyde in phosphate-buffered saline (PBS) for 1h at 4G, washed in PBS, and fixed in 1% osmium tetroxide in PBS for 1h. They were dehydrated in ascending series of graded ethyl alcohols, then in acetone. Each sample was infiltrated with the resin before being embedded in epoxy resin and polymerized for 72h. Semi-thin sections of about 0.5 to 1 μm were obtained and colored with Toluidine blue before being examined via a light microscope with an associated digital camera, hooked to a computer for image processing and editing (Leica DC 300). Ultra thin sections of about 60/90 nm were contrasted with heavy metals (uranyl acetate and lead citrate) and were examined using a Jeol 1010 transmission electron microscope at an accelerated voltage of 80kV. Images were photographed on digital images Gatan Digital Micrograph: brure Erlangen 500w: camera and edited by Image J and Microsoft Power Point.

### Human fetal single cell transcriptome analysis

Dataset GSE112570 of Single Cell RNA-Sequencing allow to explore cellular heterogeneity of human kidney cortical nephrogenic niche [8]. Experiments were performed with technology 10X Genomics single-cell RNA sequencing on two human kidney samples (17 weeks) respectively indexed in Gene Expression Omnibus (GEO) database: GSM3073088, GSM3073089. Molecular index was realized: Chromium Single Cell 3’ v2 single cell RNA-Seq of poly A selected mRNA kit (10X Genomics) and Sequencing was processed on NextSeq 500 (Illumina). Bioinformatics base call by bcl2fastq v. 2.17; reads were mapped using STAR 2.5.1b (Genome: GRCh37) and count tables were generated using the Cell Ranger software version 1.3.1. Downstream bioinformatics single cell transcriptome analyses were performed in R software version 3.4.3. Digital matrix were built with both 10X MTX files and merged in Seurat R-package version 2.3.0 [5] with package dependencies of matrix version 1.2-12, cowplot 0.9.2 and ggplot2 version 2.2.1 [21]. Batch correction was performed with canonical correlation on thirty dimensions before mathematical dimension reduction with tSNE algorithm. Also, dplyr library version 0.7.4 was used to generate intermediate table of best genes by cluster. Bioinformatics code to perform these single cell analyses was deposed at the following web address: https://github.com/cdesterke/hsckidney/.

### Kidney organoid microarray analysis

Microarray Clariom S human was done on process total RNA from human WT iPSC and its derived kidney organoids in duplicates [9]. Expression matrix was built with CEL files generated on Affymetrix Station and normalized by RMA method with TAC version 4.0 software (Appliedbiosystems) (Irizarry et al., 2003). Differential expressed genes were estimated with linear models for microarray data (LIMMA) algorithm by using a false discovery rate threshold less 5 percent [22]. Functional enrichment analysis on differential expressed genes was performed on WikiPathway database.

## Author Contributions

JWH and AGT conceived, designed, analyzed data and wrote the manuscript. JWH performed all organoids experiments and performed confocal laser scanning microscopy with analysis. JLD performed TEM, CD analyzed bioinformatics data, ABG, FG, AGT analyzed data and supervised the project. JWH, CD and AGT wrote the paper.

## Supplemental Information

**Fig. S1.**
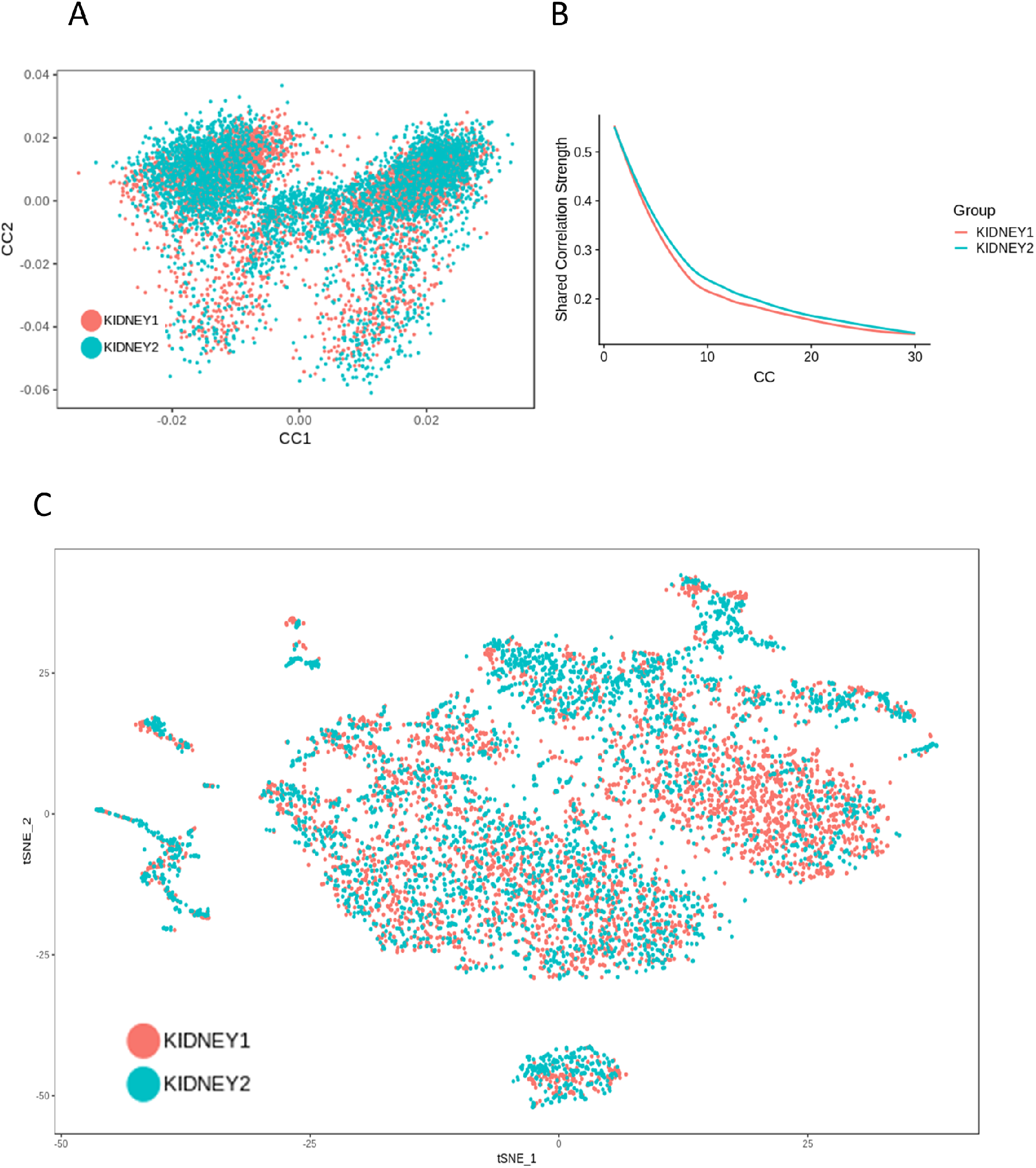
Canonical correlation with batch correction between the two foetal kidneys process in single cell transcriptome. **(a)** Factorial map showing superposition of the cells from the respective kidney during the canonical correlation. **(b)** Profile of the shared correlation strength between the respective kidneys (30 dimensions); tSNE plot post canonical correlation on the merged 2 kidneys. **(c)** t-SNE dimension reduction of single cell transcriptome from human fetal kidney after batch correction, cells from each kidney (respectively green and red) are plotted with t-SNE dimension reduction algorithm and their distribution in the map confirmed a good batch correction during the analyses (canonical correlation with Seurat R-package).

**Fig. S2.**
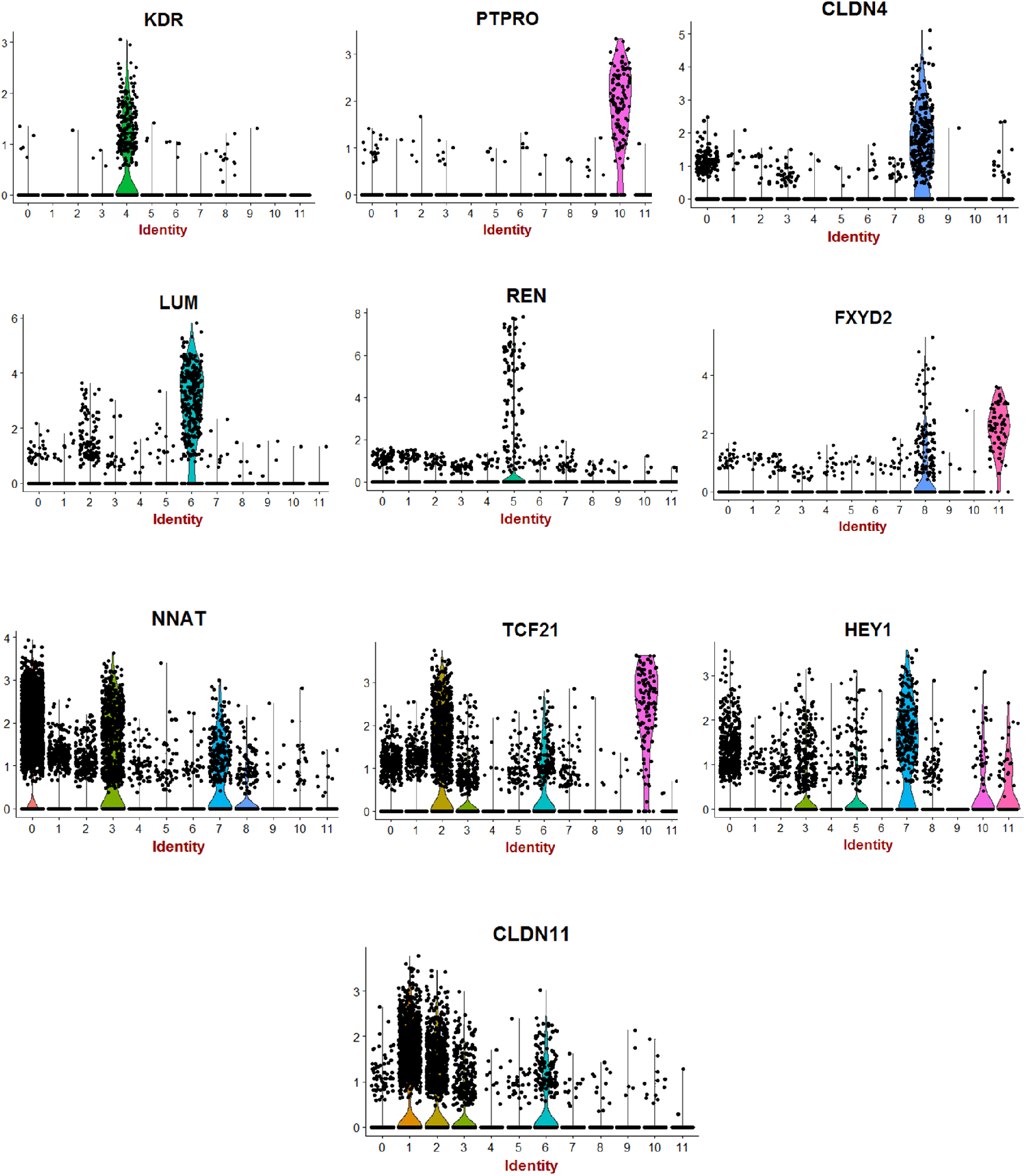
Violinplot of cluster markers found in single cell sequencing after merging the fetal cortex of two human kidneys.

**Table S1-.**
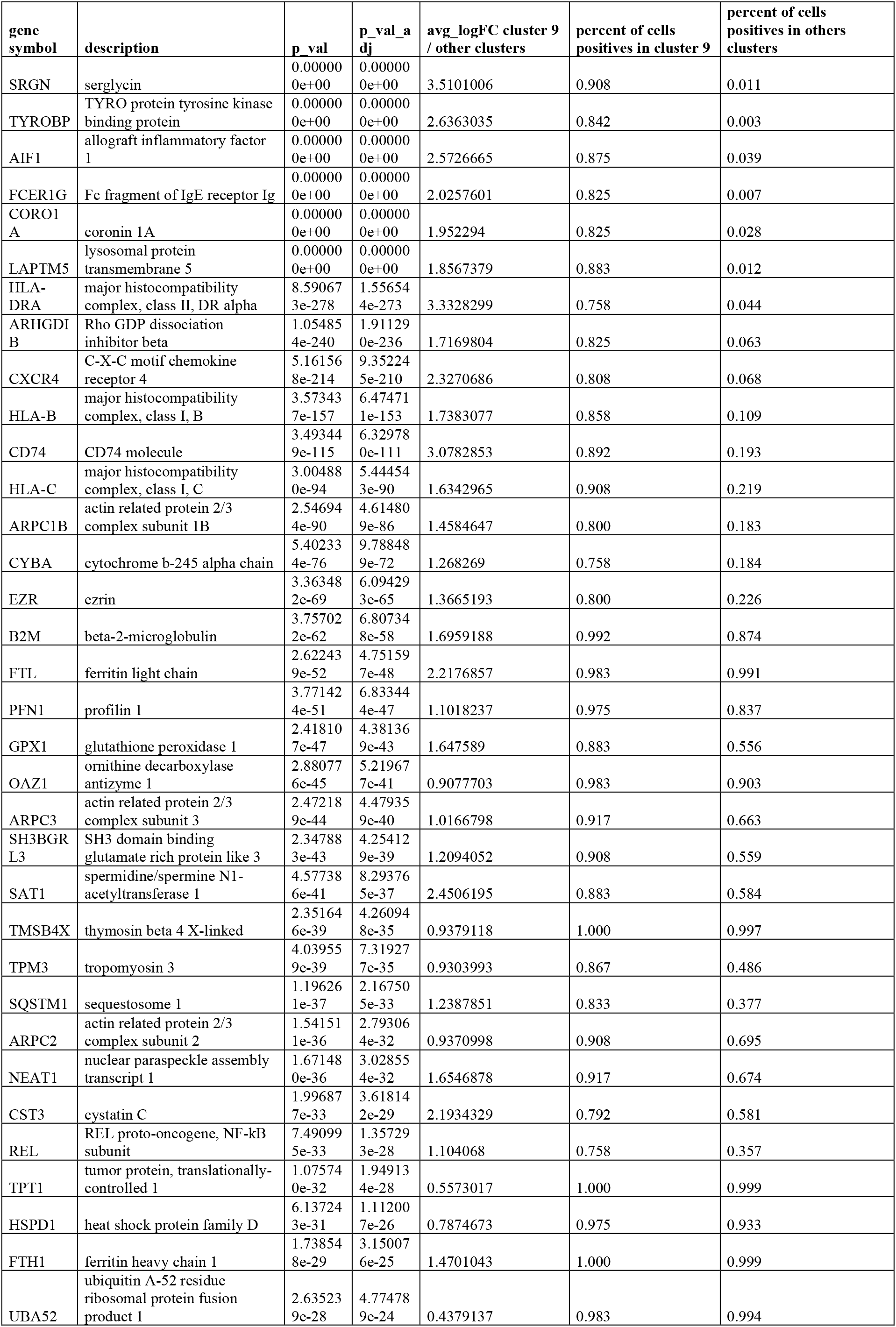
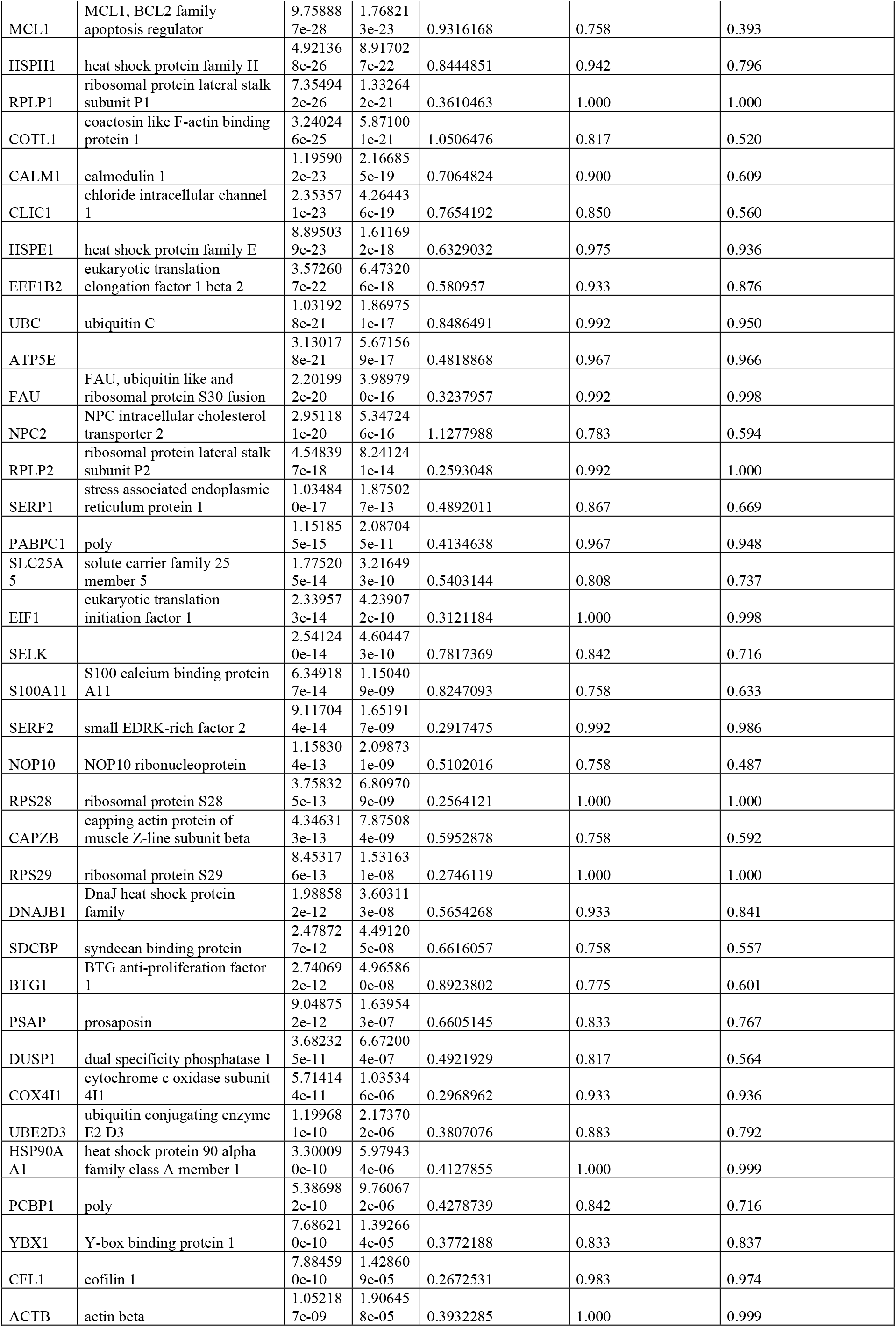
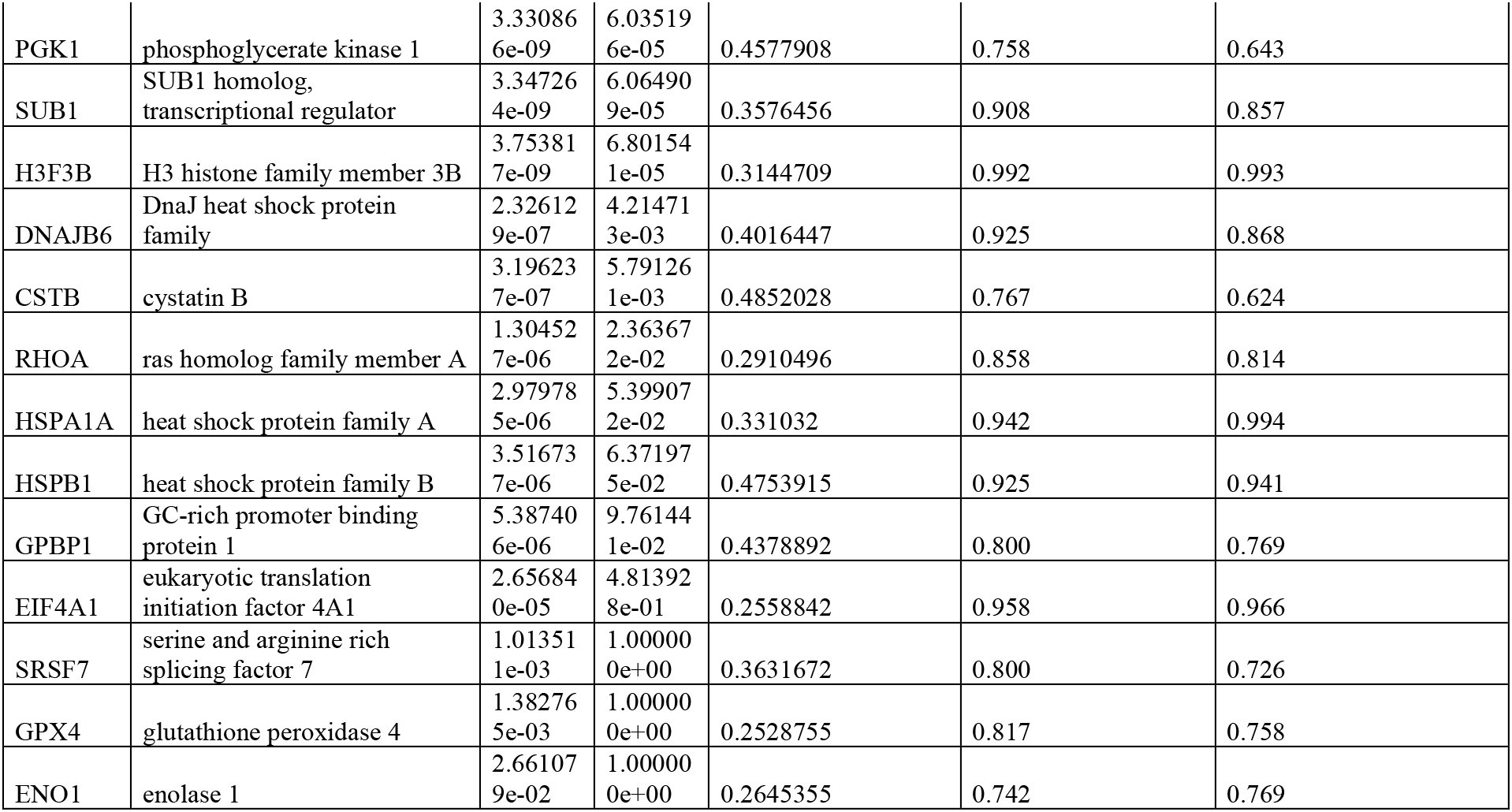
Table presenting differential expressed genes found in cluster 9 as compared to others clusters in human fetal kidney cortex.

